# Testis organ culture system capable of evaluating testicular toxicity

**DOI:** 10.1101/2023.12.01.569566

**Authors:** Kiyoshi Hashimoto, Hiroshi Arakawa, Rikako Imamura, Takuya Nishimura, Satoshi Kitajima, Takuya Sato, Kazuhide Makiyama, Takehiko Ogawa, Satoshi Yokota

## Abstract

Successful treatment of pediatric cancers often results in long-term health complications, including potential effects on fertility. Therefore, assessing the male reproductive toxicity of anti-cancer drug treatments and the potential for recovery is of paramount importance. However, *in vivo* evaluations are time-intensive and require large numbers of animals. To overcome these constraints, we utilized an innovative organ culture system that supports long-term spermatogenesis by placing the testis tissue between a base agarose gel and a polydimethylsiloxane ceiling, effectively mirroring the *in vivo* testicular environment. The present study aimed to determine the efficacy of this organ culture system for accurately assessing testicular toxicity induced by cisplatin, using acrosin-green fluorescent protein (GFP) transgenic neonatal mouse testes. The testis fragments were treated with different concentrations of cisplatin-containing medium for 24 hours and incubated in fresh medium for up to 70 days. The changes in tissue volume and GFP fluorescence over time were evaluated to monitor the progression of spermatogenesis, in addition to the corresponding histopathology. Cisplatin treatment caused tissue volume shrinkage and reduced GFP fluorescence in a concentration-dependent manner. Recovery from testicular toxicity was also dependent on the concentration of cisplatin received. The results demonstrated that this novel *in vitro* system can be a faithful replacement for animal experiments to assess the testicular toxicity of anti-cancer drugs and their reversibility, providing a useful method for drug development.

**Short Abstract:** Assessing the male reproductive toxicity of anti-cancer drugs and the potential for recovery is of paramount importance, however, *in vivo* evaluations are time-intensive and require large numbers of animals. We utilized an innovative organ culture system that mirrors the *in vivo* testicular environment to determine its efficacy in accurately assessing testicular toxicity induced by cisplatin. The results demonstrate that this system can be a faithful replacement for animal experiments to assess the testicular toxicity of anti-cancer drugs and their reversibility.

## 1. Introduction

In pharmaceuticals, it is recommended that the evaluation of testicular toxicity should be assessed in general toxicity tests through histopathological evaluation and by male reproductive toxicity tests (International Conference on Harmonization [ICH] Harmonised Guideline-S5(R3), (2020), ICH-M3(R2), (2009)). Owing to the urgency of delivering candidate treatments to patients, an assessment of the effects of anti-cancer drugs on male fertility is often not considered essential; although histopathological evaluation in general toxicity tests is required (ICH-S9, (2009)). Male reproductive toxicity tests require a considerable number of animals, a substantial amount of time, and a significant amount of compounds for testing. Currently, there are no existing guidelines that clearly indicate alternative methods as replacements for animal testing for the evaluation of male reproductive toxicity, however, *in vitro* evaluation systems are expected to supplement existing *in vivo* reproductive toxicity tests for drug discovery.

We previously developed a testis organ culture system in mice that successfully produced mature sperm from undifferentiated spermatogonia by mimicking *in vivo* intratesticular conditions (Sato et al, (2011), Sato et al, (2011), Sanjo et al, (2018)). This method used agarose gel as a platform to place the testis tissue at the gas-liquid interphase. However, the tissue inevitably contained a degenerative or necrotic region in its central area owing to a shortage of oxygen and nutrients (Sato et al, (2011), Hirano et al, (2022)), and the spermatogenic efficiency was not necessarily sufficient for use in toxicity testing (Chapin et al, (2016)). We then developed a new organ culture method using a chip made of polydimethylsiloxane (PDMS) to cover the tissue and press it thin and flat on the agarose gel. This PDMS-ceiling (PC) method, in which the oxygen concentration is lowered from ambient 20% to 15% in an incubator, has been proven to increase spermatogenic efficiency (Kojima et al, (2018), Komeya et al, (2019), Feng et al, (2023)). In addition, changes in tissue volume during cultivation could be measured by analyzing the horizontal projection area of the tissue (Hashimoto et al, (2023)).

Cisplatin is a chemotherapeutic agent frequently used to treat numerous solid tumors in pediatric oncology (World Health Organization, (2021)). In the present study, we investigated the ability of the PC testis organ culture system to evaluate the effect of clinically relevant cisplatin treatment on testicular toxicity and the time necessary for recovery. This novel alternative method addresses the problem of recovery from chemotherapy-induced testicular toxicity and potentially provides crucial information for patients concerned about fertility when considering anti-cancer medications.

## 2. Materials and methods

### 2.1 Animals

Acrosin (Acr)-green fluorescent protein (GFP)/ H3.3-mCherry double transgenic mice (Hashimoto et al, (2023)) were used in this study. These mice were generated by crossing a female Acr-GFP homogeneous mouse (Nakanishi et al, (1999)) with a male H3.3-mCherry homogeneous mouse (Makino et al, (2014)). Because both lines have been occasionally crossed with wild-type ICR or C57BL/6 mice for maintenance, their genetic backgrounds were a mixture of ICR and C57BL/6. Seven-day-old mice were used as the source of the testis tissue. We used seven mice and collected their tissues for histological evaluation at 7, 14, and 35 days after cisplatin exposure. Therefore, the number of tissue samples for the measurement of GFP fluorescence was n = 14 (ED7), n = 11 (ED14), n = 8 (ED21–35), and n = 4 (ED42–). Mice were housed in specific pathogen-free, air-conditioned rooms at 24 ± 1 °C and 55 ± 5% humidity, with a 13-hour light/11-hour dark cycle, and were fed commercially available hard pellets (MF; Oriental Yeast) ad libitum. Drinking water was acidified to pH 2.8–3.0 by HCl. All animal experiments conformed to the Guide for the Care and Use of Laboratory Animals and were approved by the Institutional Committee of Laboratory Animal Experimentation (Animal Research Center of Yokohama City University, Yokohama, Japan; protocol no. F–A–23–012).

### 2.2 Culture media and reagents

We dissolved α–modified Eagle Minimum Essential Medium powder (αMEM; 12000-022; Thermo Fisher Scientific Waltham, MA, USA) in Milli-Q purified water to twice the intended final concentration before adding an appropriate amount of AlbuMAX™ (11020–021; Thermo Fisher Scientific) to give a final concentration of 40 mg/mL. A 7% NaHCO_3_ solution was then added (0.026 mL/L) to achieve a final concentration of 1.82 g/L (0.0182 g for 10 mL of medium). Antibiotic-antimycotic (15240062; Thermo Fisher Scientific) was added to give a final concentration of 100 IU/ml for penicillin, 100 μg/mL for streptomycin, and 250 ng/mL for amphotericin. Finally, Milli-Q water was added to the medium to achieve the required volume for 1 x αMEM. The medium was sterilized with a 0.22 *μ*m Millipore filter (Burlington, MA, USA) and stored at 4 °C. In the testicular toxicity experiment, cis-diaminedichloroplatinum (cisplatin; P4394; Sigma Aldrich, St. Louis, MO, USA) dissolved in αMEM was added to the culture medium at the concentrations indicated in section 2.4. The actual cisplatin concentration was measured using liquid chromatography–tandem mass spectrometry (LC-MS/MS), as described in section 2.6.

### 2.3 PC chip production

The PDMS prepolymer and curing reagent (Silpot 184; Dow Corning, Midland, MI, USA) were mixed in a 12:1 weight ratio. The mixture was poured over a mold master, placed in a vacuum chamber for degassing, and moved to an oven at 72 °C for 1.5 h for curing. After cooling, the solidified PDMS was peeled off from the master, and this PDMS disk was diced into individual 10 mm^2^ chips using a cutter knife. The mold master was produced as previously reported, using conventional photolithography and soft lithography techniques (Duffy et a, (1998)). Each chip had a dent of 8 mm^2^ and 160 μm deep on one side, which served as a space for accommodating the tissue fragment.

### 2.4 Culture method

Agarose gel blocks (1.5% w/v) were prepared by dissolving agarose powder (346-00072, Dojindo, Japan) in purified Milli-Q water (Millipore) and autoclaving. During cooling, 33 mL of the agarose solution was poured into 10 cm dishes to form a 5 mm thick gel. The gel was then cut into 10 mm square pieces to be used as platforms for testis tissue placement. The gels were placed in a 12-well culture plate and submerged twice in the culture medium for more than 6 h, with a medium change in between. After removing the second medium, 0.5 ml of fresh medium was added to each well to half soak the gels.

The testes were decapsulated and divided into six pieces of equal pieces using forceps. The tissue fragments were then placed on blocks of agarose gel in a 12-well culture plate (665180; Greiner Bio–One, Austria). A single tissue was placed on a gel and a PC chip with a dent depth of 160 μm was placed over the tissue (Kojima et al, (2018), Komeya et al, (2019)) (Fig. 1A). The culture plates were placed in an incubator at 34 °C with 15% oxygen and 5% carbon dioxide. The medium was changed once a week and was poured up to half the height of the agarose gel (approximately 0.5 mL/well). For cisplatin treatment, a series of culture media were prepared with specified concentrations of cisplatin: 0, 0.4, 1, 4, 12, and 40 μg/mL, designated as cis-0, cis-0.4, cis-1, cis-4, cis-12, and cis-40, respectively. The original medium was replaced with the new media with cisplatin, and testis tissues were exposed to the cisplatin for 24 h. The media containing cisplatin were then replaced with αMEM containing AlbuMAX medium, followed by a series of three washes at two-hour intervals, using fresh culture medium each time to remove any residual cisplatin.

**Figure. 1.**
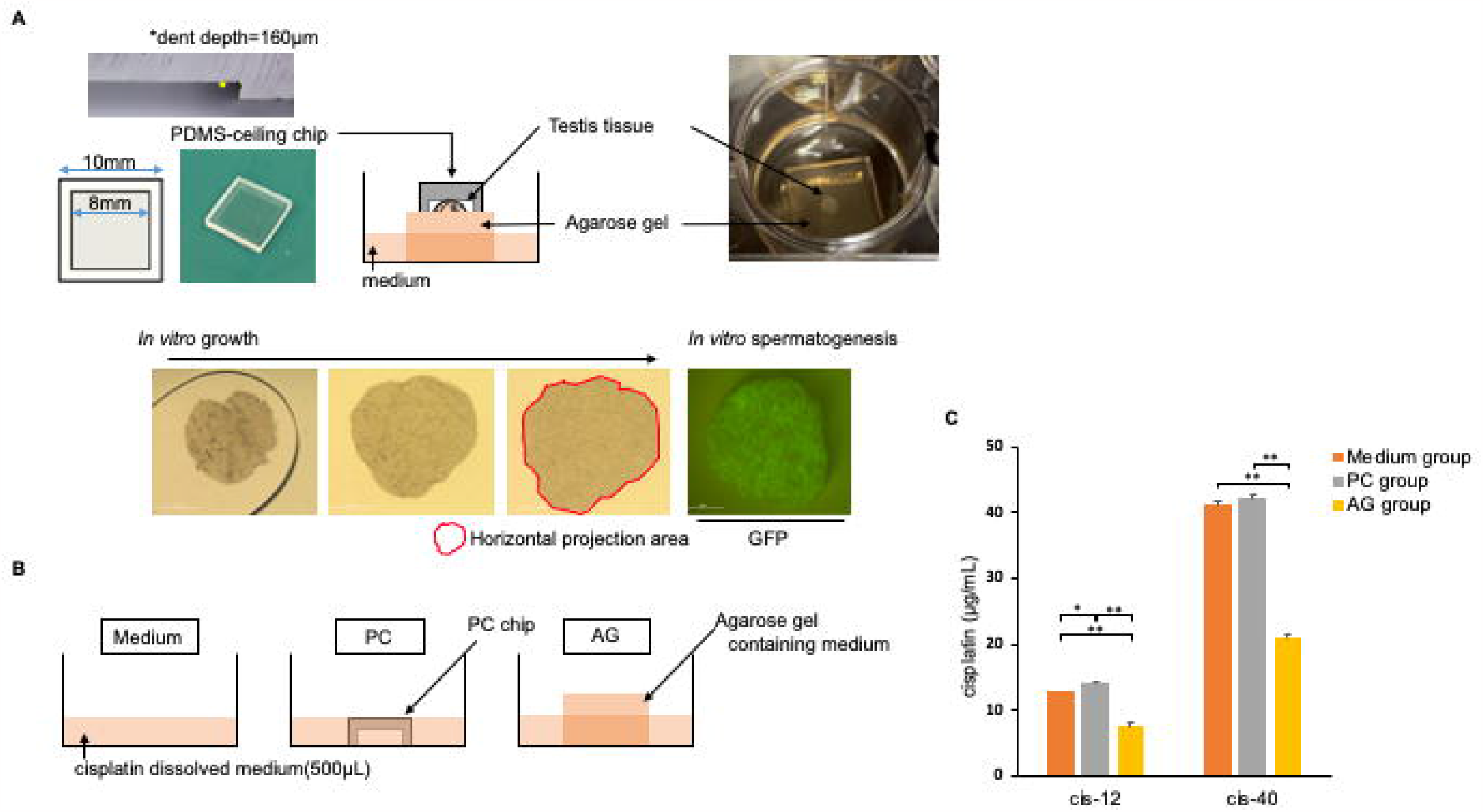
Polydimethylsiloxane-ceiling method (PC method). **A**. Schematic presentation of the PC method. The testicular tissue is flattened inside the narrow dent space, and allows the monitoring tissue growth and spermatogenesis *in vitro* over time without interruption of culture. **B**. Experimental setup to investigate the effects of the experimental components. Medium, containing only cisplatin dissolved medium; PC, containing cisplatin dissolved medium with a PC chip; and AG, containing cisplatin dissolved medium and medium-filled agarose gel. **C**. Summary graph representing the change in the cisplatin concentration after 24 hours in the Medium, PC, and AG groups. Error bars indicate SEM. Statistical analysis by Tukey HSD test. * *p* < 0.05, ** *p* < 0.01 (compared with cis-0). cis-12 and cis-40: cisplatin concentration, 12 and 40 *μ*g/mL, respectively.

### 2.5 Identification of the effects of the *in vitro* culture system components

To determine the separate effects of the PC chip and the agarose gel, we conducted an additional experiment with three distinct experimental setups for the cis-12 and cis-40 concentrations. The setups were: Medium group, with only cisplatin-infused medium in the wells; PC group, comprising cisplatin-infused medium with PC chips; and AG group, which contained cisplatin-infused medium and a fresh agarose gel (Fig. 1B).

### 2.6 Measurement of cisplatin concentrations

To accurately determine the cisplatin concentration, a portion of the medium was collected 24 h after the addition of each cisplatin concentration. The cisplatin concentration was measured by LC-MS/MS after derivatization (Shaik et al, (2016), Arakawa et al, (2022)) with slight modifications. Briefly, 10 *μ*L of the supernatant was mixed with 10 *μ*L of 15 mg/mL diethyldithiocarbamate (FUJIFILM Wako Pure Chemical Corporation, Osaka, Japan) in 0.1 mol/L NaOH and 65 *μ*L of methanol and incubated at 40 °C for 30 min. After centrifugation at 21,600 ×*g* for 10 min at 4 °C, the resulting supernatant was collected and analyzed using LC-MS/MS. The derivatized cisplatin was measured using a triple quadrupole mass spectrometer LCMS8050 (Shimadzu, Kyoto, Japan) coupled to an LC30A system (Shimadzu). Chromatography was performed on a CAPCELL PAK C18 MG III (ID 2.0 mm × 50 mm; Shiseido Co. Ltd., Tokyo, Japan) at 40 °C. The mobile phase was composed of 0.1% formic acid in water (solvent A) and 0.1% formic acid in acetonitrile (solvent B) using a flow rate of 0.3 mL/min, and the injection volume was 10 *μ*L. The mobile phase gradient elution started at 5% solvent B for 0.5 min, followed by a linear gradient to 75% solvent B for 1 min, a linear gradient to 90% solvent B for 1.5 min, a linear gradient to 95% solvent B for 2.5 min, maintained at 95% solvent for 1.5 min, and then returned to the initial conditions for 1 min. Electrospray positive ionization was used, and mass transitions were monitored at m/z 639 > 491. LabSolutions software (Shimadzu) was used for data manipulation.

### 2.7 Observations of GFP fluorescence in the testes

Cultured tissues were observed at least once a week under a stereomicroscope equipped with an excitation light for GFP (Leica M205 FA; Leica, Germany). For semi-quantitative evaluation of Acr-GFP expression, cultured testis tissues were classified into six grades according to the percentage of GFP-positive areas: 0, 0%; 1, 1–20%; 2, 21–40%; 3, 41– 60%; 4, 61–80%; and 5, 81–100% (Supplemental Fig. 1). This GFP grading scale has previously been demonstrated to accurately correspond to spermatogenic progression (Sato et al, (2011), Komeya et al, (2016)). The horizontal projection area of the tissue was measured using Fiji, an open-source platform for biological image analysis (Schindelin et al, (2012)). The tissue volume was calculated by multiplying the measured horizontal projection area by the depth of the PC chip dent. The relative tissue area and volume of each cultured tissue were compared with those measured on culture day 0.

### 2.8 Histological examinations of the testes

For histological examination, specimens were fixed with Bouin’s fixative and embedded in paraffin. Each specimen was stained with hematoxylin and periodic acid Schiff (PAS) (MUTO PURE CHEMICALS). The differentiation stage of spermatogenic germ cells was determined based on morphological characteristics, specifically the chromatin pattern within the nucleus and the acrosomal staining profile. To investigate the differentiation of germ cells in cultured tissues after cisplatin treatment, we measured the percentage of seminiferous tubules with pachytene spermatocytes (%STp) and those with haploid cells (%STh) out of the total number of seminiferous tubule cross sections in the cultured tissue based on the PAS-stained images.

### 2.9 Statistical analysis

Results are presented as mean ± SE. Statistical analysis was performed using one-way analysis of variance, followed by the Tukey – Kramer HSD test. A p-value of *p* < 0.05 was considered statistically significant.

## 3. Results

### 3.1 Effects of experimental equipment on cisplatin concentration

The actual concentrations of cisplatin to which the tissue was exposed were thought to vary owing to factors such as the presence of the agarose gel and the PC chip. Therefore, the cisplatin concentration in the media after a 24-hour period in culture wells containing both the agarose gel and the PC chip was quantified. The LC-MS/MS results showed that the initial cisplatin concentrations set at 4 *μ*g/mL, 12 *μ*g/mL, and 40 *μ*g/mL decreased to 2.44 ± 0.48 *μ*g/mL, 7.52 ± 0.54 *μ*g/mL, and 21.0 ± 1.4 *μ*g/mL, respectively; suggesting an approximate halving of the concentration within 24 hours. Concentrations of cisplatin at 1 *μ*g/mL or lower were below the LC-MS/MS detection threshold and were not quantifiable. In the Medium group, LC-MS/MS analyses revealed cisplatin levels of 12.9 ± 0.7 *μ*g/mL and 41.2 ± 2.8 *μ*g/mL for cis-12 and cis-40, respectively, and there were negligible changes observed in the PC group. This indicated that the PC chip had a minimal impact on cisplatin concentration. In contrast, in the AG group, the cisplatin levels for cis-12 and cis-40 were measured at 7.52 ± 0.54 *μ*g/mL and 21.0 ± 1.4 *μ*g/mL, respectively; suggesting that the cisplatin concentration was halved (Fig. 1C). This reduction aligns with the premise that the medium volume (500 *μ*L) and the agarose gel dimensions (10 mm × 10 mm × 5 mm) would result in the dilution of cisplatin owing to the water content of the agarose gel. The LC-MS/MS measured concentrations of cisplatin were used to establish exposure levels in subsequent experiments.

### 3.2 Effects of cisplatin treatment on testicular tissue volume

The present study was performed as shown in the experimental flow (Fig.2A). The bright field for the treated testicular tissues are shown in Fig. 2B. During this experimental period, we observed changes in tissue volume every week (Fig. 2B). In the control group (cis-0), the tissue volume gradually increased, with an approximately three-fold increase at 49 days after exposure (ED49) compared that at ED7 (Fig. 2B, C). The volume in cis-0.4 and cis-1 groups also increased, similar to that of the control group (Fig. 2B, C). In contrast, the cis-4 and higher concentration groups showed significant tissue shrinkage compared with the control until ED14; after which the cis-4 and cis-12 groups showed a rapid increase, and there were no significant differences compared with the control from ED21 onward. However, tissue volume in the cis-40 group did not increase during the observation period (Fig. 2B, C).

**Figure. 2.**
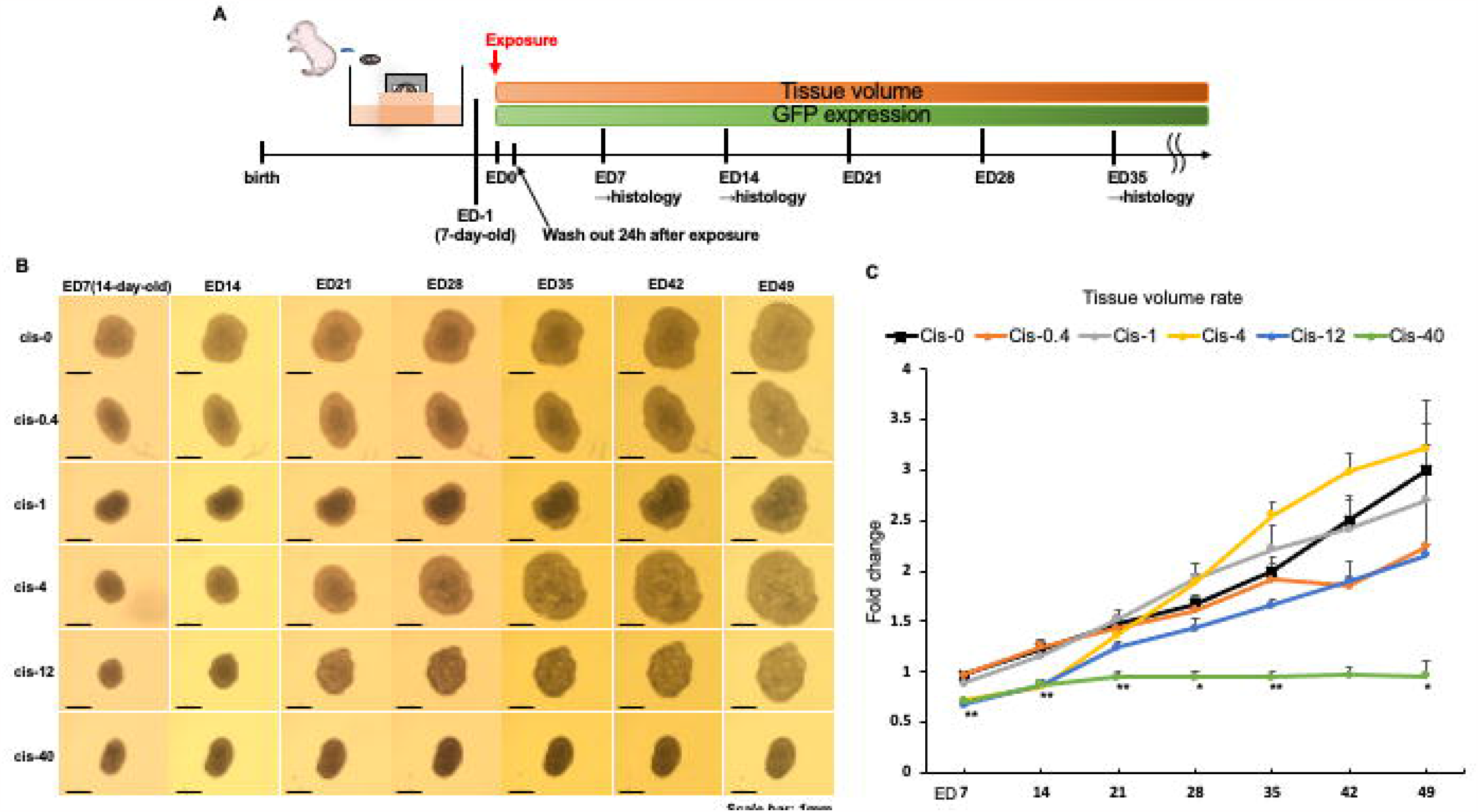
Testicular tissue changes after cisplatin exposure. **A**. Experimental schema. Normal medium was replaced with cisplatin dissolved medium (cis-0, cis-0.4, cis-1, cis-4, cis-12, and cis-40) and cultured for 24 hours. The medium was then replaced with normal medium. Subsequently, the changes in tissue volume and transition of GFP expression were observed every week. **B**. Stereomicroscopic view of cultured tissues. Scale bar: 1 mm. **C**. Relative tissue volume compared to the volume seven days after exposure (ED7). No significant differences were observed in groups below cis-1. Cis-4 or higher groups showed significant tissue shrinkage compared with cis-0 until ED14. Cis-40 showed no tissue increase during the observation period. Error bars indicate SEM. Statistical analysis by Tukey HSD test. * *p* < 0.05, ** *p* < 0.01 (compared with cis-0) cis-0, cis-0.4, cis-1, cis-4, cis-12, and cis-40: cisplatin concentration, 0, 0.4, 1, 4, 12, and 40 *μ*g/mL, respectively.

### 3.3 The effects of cisplatin treatment on spermatogenesis

The germ cells of Acr-GFP mice expressed GFP in the cytoplasm as they progressed to the pachytene stage (stage IV), allowing the evaluation of germ cell differentiation over time without interrupting cultivation (Supplemental Fig. 1A, B). For quantitative evaluation, the percentage of GFP expression in the cultured tissue area was graded into six levels: grades 0–5 (Supplemental Fig. 1B). The GFP expression observed in the tissues is shown in Fig. 3A and B. At the time of cisplatin treatment (ED0), GFP expression was not observed in any of the tissues as they had only been cultivated for a period equivalent to the maturation stage of eight-day-old tissues. At ED7, GFP expression was almost equal among the control, cis-0.4, cis-1, and cis-4 groups, however, the cis-12 and cis-40 groups showed very little GFP expression. At ED14, the GFP grade in the cis-0.4 and cis-1 groups remained almost equivalent to that of the control. Interestingly, cis-4 showed a significant decrease in GFP expression levels at ED14, followed by a rapid recovery to a level comparable to that of the control at ED35. In cis-12, a partial recovery of GFP expression was observed around ED35, after which GFP expression gradually improved until ED70 (Supplemental Fig. 2). However, no GFP expression was detected in cis-40 until ED70 (Supplemental Fig. 2).

**Figure. 3.**
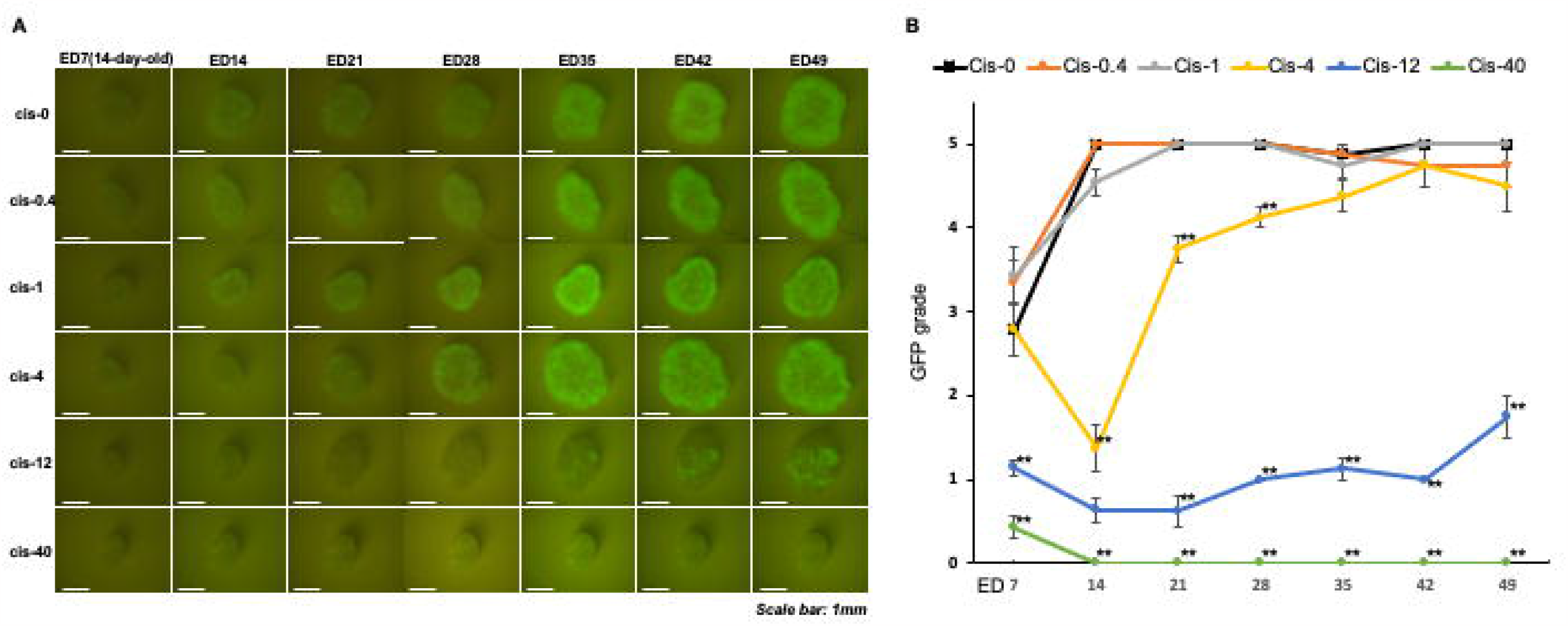
Spermatogenesis after cisplatin tratment. **A**. Stereomicroscopic image of cultured tissues under excitation showing GFP expression in the seminiferous tubules. Scale bar: 1 mm. **B**. Summary graph representing GFP grade in each cisplatin concentration. No significant difference was found in groups below cis-1. Error bars indicate SEM. Statistical analysis by Tukey HSD test. ** *p* < 0.01 (compared with cis-0) cis-0, cis-0.4, cis-1, cis-4, cis-12, and cis-40: cisplatin concentration, 0, 0.4, 1, 4, 12, and 40 *μ*g/mL, respectively.

### 3.4 Histological results after cisplatin exposure

Histological examination of the effects of cisplatin treatment using PAS staining was performed at ED7, ED14, and ED35 (Fig. 4A); and the percentage of STp (Fig. 4B) and STh determined (Fig. 4C). The results showed that the %STp in the cis-0.4 and cis-1 groups was not significantly different from that in the control at any time point. In contrast, there was a significant decrease in %STp at ED14 in the cis-4 group, which recovered to a level almost equivalent to that of the control at ED35. In the cis-12 group, a significant decrease in %STp was found at ED7, with almost no meiotic germ cells observed at ED14. Subsequently, the damage recovered by ED35, showing meiotic germ cells in some seminiferous tubules. In the cis-40 group, histological analysis revealed extensive germ cell loss and degeneration at ED7, and only Sertoli cells remained in most of the seminiferous tubules at ED14 and ED35. In the control and cis-0.4 groups, haploid cells were found in over 40% of the seminiferous tubules in cultured tissue (cis-0: 44.9%, cis-0.4: 42.5%), and there was no significant difference between the groups. The %STh was significantly lower in the cis-1 and higher concentration groups compared with cis-0.

**Figure. 4.**
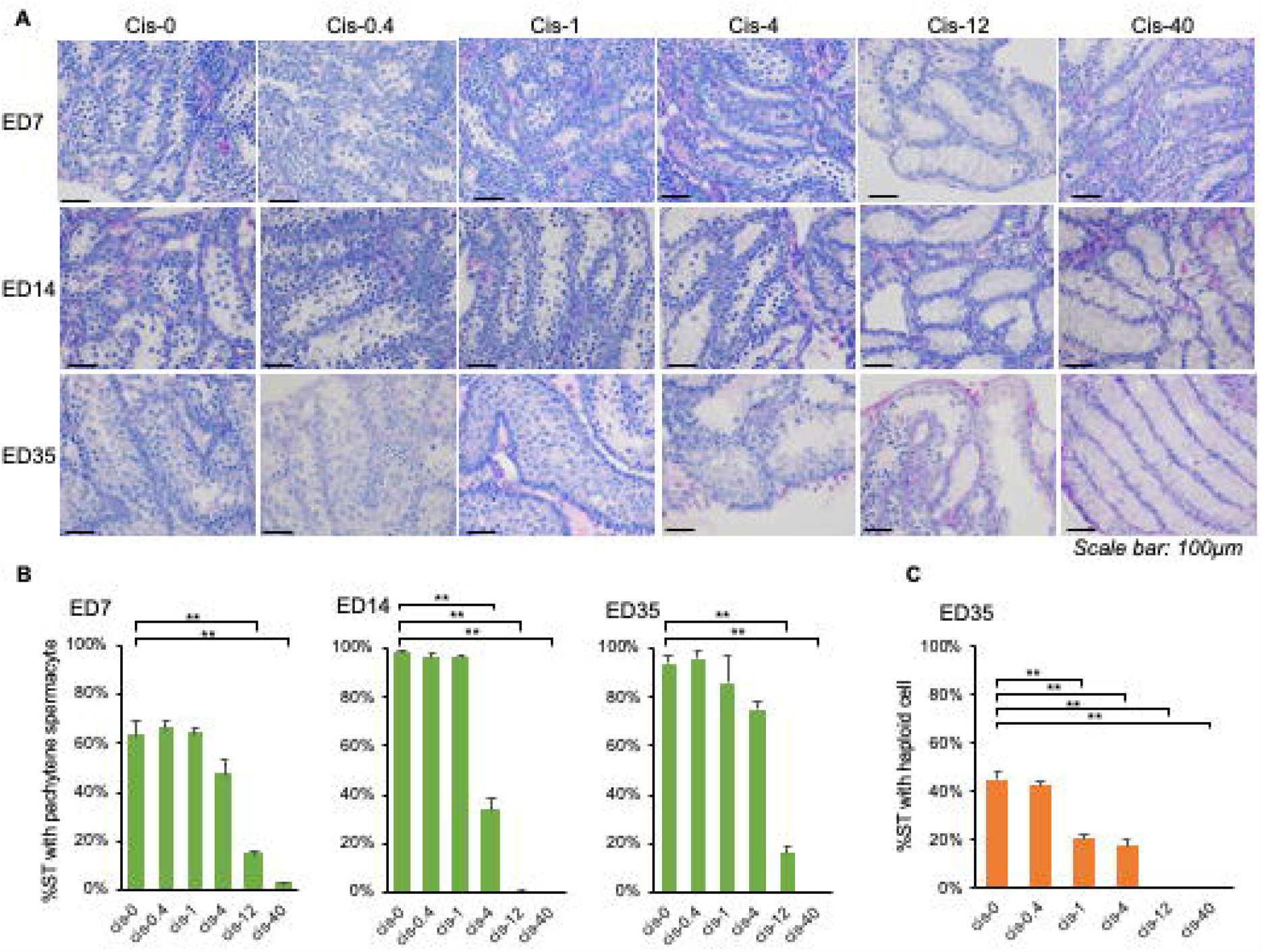
Histological findings after cisplatin treatment. **A**. Periodic acid Schiff staining for the cultured tissue at ED7, ED14, and ED35. Scale bar: 100 μm. **B**. %STp at each cisplatin concentration at ED7, ED14, and ED35. **C**. %STh at each cisplatin concentration at ED35. %STh in cis-1 was significantly lower than that in cis-0, although the tissue volume changes and GFP grade were not significantly different. There were three samples in each group. Error bars indicate SEM. Statistical analysis by Tukey HSD test. ** *p* < 0.01 (compared with cis-0) %STp: the percentage of seminiferous tubules with pachytene spermatocytes out of the total number of seminiferous tubule cross–sections, %STh: the percentage of seminiferous tubules with haploid cell out of the total number of seminiferous tubule cross–sections. Error bars indicate SEM. Statistical analysis by Tukey HSD test. ** *p* < 0.01 (compared with cis-0) cis-0, cis-0.4, cis-1, cis-4, cis-12, and cis-40: cisplatin concentration, 0, 0.4, 1, 4, 12, and 40 *μ*g/mL, respectively.

## 4. Discussion

Taking the 3Rs of animal experiments into consideration, it would be advantageous to use as few animals as possible in toxicity evaluations. However, the development of a simple and robust alternative screening method for evaluating male reproductive toxicity, such as *in vitro* spermatogenesis, is slow. The present study demonstrated the effectiveness of the Acr-GFP transgenic mouse pre-pubertal testis organ culture system (Sato et al, (2011), Sanjo et al, (2018), Sato et al, (2011), Kojima et al, (2018), Komeya et al, (2019), Yokonishi et al, (2014)) to evaluate testicular toxicity after cisplatin exposure and recovery. In Acr-GFP transgenic mice, GFP is expressed in spermatocytes at the mid-pachytene stage of meiosis onward (Ventela et al, (2000)). Using the testes of these mice allows the spermatogenic progression at or beyond this stage to be monitored sequentially while maintaining the culture experiment. This permits the sequential evaluation of spermatogenic impairment and recovery in real time by observation of GFP expression. In addition, sequential measurements of the tissue volume reflected the testicular toxicity of cisplatin. These evaluation systems could serve as a general means to identify male reproductive toxicity for any chosen chemical.

Survival rates for childhood cancers have significantly improved in recent decades, with the current five-year survival rate for most childhood cancers now exceeding 85% (Miller et al, (2022)). Among the various treatment modalities, cisplatin-based chemotherapy continues to be considered the mainstay, and cisplatin remains one of the most commonly used drugs for treating various solid tumors in pediatric patients (World Health Organization, (2021)). However, cisplatin-based chemotherapy induces spermatogenic impairment and fertility problems as late adverse events (Qi et al, (2019), Tharmalingam et al, (2020)). Therefore, for pediatric cancer patients, information about the gonadal toxicity of each anti-cancer drug, particularly whether the damage caused by them is temporary or permanent, is crucial to maximize the effect of these drugs on the disease while minimizing the damage to the testis and future fertility. To date, no studies have examined the direct impact of cisplatin on the testes of children owing to the scarcity of testicular samples collected before and after chemotherapy. Although some animal studies have reported that cisplatin treatment causes testicular toxicity (Oshio et al, (1990), Wang et al, (2018), Zhang et al, (2001), Seaman et al, (2003), Sawhney et al, (2005), Harman and Richburg, (2014)), its reversibility has rarely been evaluated. Therefore, this study used an organ culture system to investigate this subject. The results showed that the clinically-relevant cis-4 group, but not the lower groups, showed testicular toxicity but recovered within one cycle of the seminiferous epithelium in mice. A possible therapeutic range for the concentration of cisplatin in the serum of most cancer patients has been reported as 1–4 μg/mL (Dominici et al, (1989), Erdlenbruch et al, (2001), Panteix et al, (2002), Ikeda et al, (1998), Charlier et al, (2004)). Our results suggest that contraception periods should be set to at least one cycle of seminiferous epithelium to avoid male infertility in the case of clinically-relevant administration of cisplatin. Cisplatin administration results in the formation of cisplatin–DNA intra- and interstrand adducts. Failure to repair these lesions initiates apoptosis (Wang et al, (2018)), leading to substantial elimination of rapidly dividing cell populations (Kelland, (2007)). Animal experiments have revealed that a single dose of cisplatin induces apoptosis in male germ cells, leading to decreased spermatogenesis (Zhang et al, (2011), Seaman et al, (2003)). Our results showed that expression of GFP from pachytene spermatocytes to spermatids was decreased by cisplatin treatment, which may be due to increased apoptosis of male germ cells.

Previous studies have investigated the effects of chemicals on testicular toxicity in a testis organ culture system through histological examination (Nakamura et al, (2017), Nakamura et al, (2019), Smart et al, (2018), Lopes et al, (2020)). The results of the present study indicate that the GFP grading scale reflects the histopathological findings in the testes. Compared to any other *in vitro* culture models (Steinberger et al, (1964), Trowell, (1959), Haneji et al, (1983), Rassoulzadegan et al, (1993), Nagao, (1989), Abu et al, (2012)), our organ culture system was superior by allowing the effective progression of spermatogenesis and incubation for a longer time period thereby permitting the identification of the recovery period. In a previous study, clinically-relevant exposure levels of cisplatin, at concentrations of 0.25, 0.5 or 0.75 μg/mL, impaired spermatogenesis in neonate testis fragments under cultivation. Recovery from this damage was not definitely recognized until eight weeks after the treatment (Lopes et al, (2020)). Our method of culturing testicular fragments on agarose gel with a PC chip covering improved spermatogenic efficiency and increased its duration to over 70 days. Owing to such robust and long-lasting spermatogenesis *in vitro*, we were able to monitor the damage induced by cisplatin, as well as its recovery. Therefore, our culture system is a useful tool for the toxicological screening of anti-cancer drugs.

In most clinical settings, several anti-cancer drugs are used in combination, the so-called regimen, rather than single-agent monotherapy. Each regimen involves several doses of agents over a set number of days (Rossi and Di, (2016), Dear et al, (2013)), and it is important to note that toxicological evaluations should consider this repeated drug administration. Future studies are required to investigate the effects of multiple cycles of drug exposure on testicular toxicity and recovery. The results would allow the design and optimization of new chemotherapy regimens for each pediatric cancer patient.

PDMS is widely used in the fabrication of microfluidic devices and microphysiological systems. However, it has been reported that PDMS can absorb various chemicals from a medium (Van et al, (2017)), complicating the measurement and precise control of these chemicals. Consequently, we measured the cisplatin concentration after a 24-hour incubation in a well containing the PC chip and agarose gel. Our findings revealed no significant effect of the PC chip on the cisplatin concentration, possibly because cisplatin is water-soluble, whereas chemicals absorbed by PDMS are typically lipid-soluble (Toepke and Beebe, (2006), Auner et al, (2019)). In contrast, the presence of agarose gel halved the cisplatin concentration. This outcome is reasonable, given that the volumes of the gel and administered medium were equal. Although this may be considered a drawback of using agarose gel, the concentration of the applied chemicals can be appropriately adjusted based on the volume of the agarose gel and medium used in each experimental setup.

In conclusion, our findings provide a basis for the continued development of an effective *in vitro* testis organ culture system with a reduced time and number of animals required. This may be useful to supplement current male reproductive toxicity testing or as an alternative model in cases where *in vivo* testing may not be feasible. Further experiments and refinements are required to understand the extent of testicular toxicity and its reversibility with respect to this new modality of chemotherapy in our culture model. We anticipate that through our new evaluation method, we can provide rapid information to support the appropriate use of each drug for patients, in the hope of maintaining their future reproductive potential.

## Author contribution

KH, TN, TO and SY wrote the manuscript. All authors discussed the results of the study. SY conceived the study. TS and TO established the organ culture method. KH, HA, and RI conducted the experiments. SK, TS, and KM supervised the study.

## Data and materials availability

All the data and materials used in this study are available in some form upon request from any of the authors to reproduce or extend the analysis.

## Acknowledgments

We thank Margaret Biswas, Ph.D., from Edanz (https://jp.edanz.com/ac), for editing the draft of this manuscript.

## Figure legends

**Supplementary Figure 1: GFP grading scale**

**A**. Mouse seminiferous epithelial cycle (green cover: the germline with GFP expression). Translucent green cover shows the germline with GFP expression after pachytene spermatocytes stage IV. **B**. GFP grading scale (PC method version). We classified into six grades based on the percentage of GFP expression area under a stereomicroscope with excitation light. Grade 0 was no GFP expression and the other grade was defined according to the percentage of GFP expression area (Grade1: 1–20%, Grade2: 21–40%, Grade3: 41–60%, Grade4: 61–80%, Grade5: 81–100%).

**Supplementary Figure 2: Tissue changes and GFP expression after ED49 in high cisplatin treatment groups**.

Stereomicroscopic views of tissue changes and GFP expression in cultured tissues after ED 49. Each cisplatin concentration was 0, 12, 40 μg/mL (cis-0, cis-12, cis-40) Scale bar: 1 mm.

