## Supplementary figures and images for "Testis organ culture system capable of evaluating testicular toxicity"

### Supplemental Figure. 1

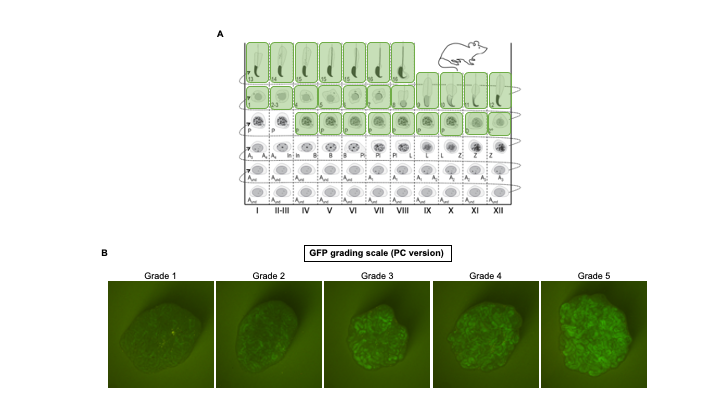

### Supplemental Figure. 2

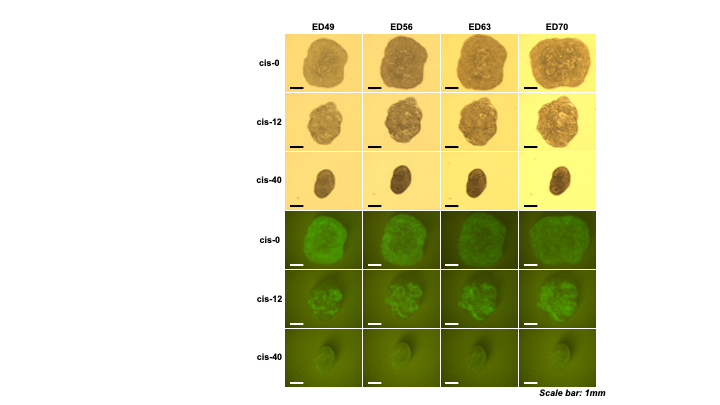
